# IK channel confers fine-tuning of rod bipolar cell excitation and synaptic transmission in the retina

**DOI:** 10.1101/2024.05.21.595126

**Authors:** Yong Soo Park, Ki-Wug Sung, In-Beom Kim

## Abstract

During retinal visual processing, rod bipolar cells (RBCs) transfer scotopic signals from rods to AII amacrine cells as second-order neurons. Elucidation of the RBC excitation/inhibition is essential for understanding the visual signal transmission. Although excitation and extrinsic inhibitory mechanisms have been studied, intrinsic inhibitory mechanisms remain unclear. We focused on RBC’s prominent K^+^ current, which exhibits voltage and Ca^2+^ dependence. We isolated and confirmed intermediate-conductance Ca^2+^-activated K^+^ channels (IK) and in RBCs using the patch-clamp method with IK inhibitors (clotrimazole and TRAM34). The regulation of the IK current primarily relies on Ca^2+^ influx via low-threshold Ca^2+^ channels during RBC excitation. It mediates RBC repolarization and oscillation, enabling fast and transient synaptic transmission to AII amacrine cells. Our findings highlight the unique role of the IK channel in RBC, suggesting that it plays a critical role in the scotopic pathway by fine-tuning RBC activity.

## Introduction

Rod bipolar cells (RBC) is the central mediator of the scotopic pathway in the retina. RBC receives a scotopic visual signal from the rod photoreceptor and transmits it to the AII amacrine cell, which contacts the cone bipolar cell terminal to convey the scotopic signal to the ganglion cell (Chun et al, 1993; Demb & Singer, 2012; Kolb & Famiglietti, 1974; Strettoi et al, 1990, 1994; Taylor & Smith, 2004). Rod bipolar cells have distinct excitatory components compared with other conventional neurons in the central nervous system (CNS). That is, it expresses metabotropic glutamate receptor 6 (mGluR6) and transient receptor potential cation channel subfamily M member 1 (TRPM1), instead of ionotropic glutamate receptors, such as α-amino-3-hydroxy-5-methyl-4-isoxazolepropionic acid (AMPA) and N-methyl-D-aspartate (NMDA) (Koike et al, 2010; Morgans et al, 2010; Vardi et al, 2000; Vardi & Morigiwa, 1997). Additionally, it expresses T-type and L-type voltage-gated Ca^2+^ channels (VGCC) without voltage-gated Na^+^ channels. These distinct characteristics generate a gradual potential for synaptic transmission rather than an action potential (Pan, 2000; Protti & Llano, 1998; Tachibana et al, 1993; Wan et al, 2008).

To understand the precise visual processing and transmission in RBCs, their inhibitory and excitatory mechanisms need to be elucidated. Therefore, inhibitory synaptic inputs to RBCs from horizontal and amacrine cells have been described in detail (Yang, 2004). Horizontal cells release the γ-aminobutyric acid (GABA) onto the dendrites of the bipolar cells in the outer plexiform layer (OPL) for feedforward inhibition and formation of the receptive field (Ströh et al, 2018; Yang & Wu, 1991). And the A17 amacrine cells release GABA to the axon terminal of the RBCs, located in the innermost layer of the inner plexiform layer (IPL), as a reciprocal feedback inhibition to control the pre-synaptic glutamate release from RBCs (Elgueta et al, 2018; Grimes et al, 2009; Grimes et al, 2015; Hartveit, 1999). Additionally, GABAergic amacrine cells synapse into RBCs, which are modulated by the dopamine pathway (Travis et al, 2018). However, there is limited information regarding the intrinsic inhibitory properties of RBCs, such as K^+^ and Cl^-^ channels, which are controlled by membrane potential or intracellular Ca^2+^. Although various types of K^+^ channels and Cl^-^ channels were characterized in the nervous system as important regulators for neural excitation (Burke & Bender, 2019; Lai & Jan, 2006; Rahmati et al, 2018), only a few were discussed in RBCs (Van Hook et al, 2019). For example, voltage-gated K^+^ channels, voltage-gated Cl^-^ channels, and Ca^2+^-activated Cl^-^ channel, anoctamin 1 (ANO1) were identified in the RBCs using electrophysiology and immunohistochemistry. However, their role in excitability has not yet been described (Enz et al, 1999; Hu & Pan, 2002; Jeon et al, 2013; Klumpp et al, 1995; Paik et al, 2020). Nevertheless, those identified channels cannot sufficiently explain the prominent K^+^ current in RBCs, and their functional role in excitability and synaptic transmission in RBCs has not been revealed. Thus, more information on K^+^ channels is required to understand the mechanisms controlling RBCs.

Ca^2+^-activated K^+^ channels are classified into three types based on their conductance, defined as large (BK), intermediate (IK), and small (SK). Those are critical for modulating the excitatory post-synaptic potential, action potential, after-hyperpolarization, and neurotransmitter release in neurons (Kshatri et al, 2018). The expression and function of Ca^2+^-activated K^+^ channels have been studied in retinal neurons. The BK channels have been found in the photoreceptors, on-bipolar cells, horizontal cells, amacrine cells, and ganglion cells in various species (Burrone & Lagnado, 1997; Ingram et al, 2020; Llobet et al, 2003; Moriondo et al, 2001; Pelucchi et al, 2008; Sun et al, 2017). Additionally, the localization of SK channels in horizontal cells, amacrine cells, and ganglion cells has been reported in rodent retinas (Clark et al, 2009; Klöcker et al, 2001; Pelucchi et al., 2008; Wang et al, 1998). These studies showed that BK and SK are mainly involved in the depolarization, spiking activity, and spontaneous oscillations of retinal neurons. However, they are not expressed in RBCs (Paik et al., 2020; Tanimoto et al, 2012). Meanwhile, IK expression in the retina has not been widely explored, except for a previous report showing IK localization in the salamander photoreceptors (Pelucchi et al., 2008). Therefore, we assume that IK can be expressed in RBCs as an intrinsic inhibitory regulator coupled with VGCC to control Ca^2+^-dependent excitation.

To test our hypothesis, we used the patch-clamp method and recorded the K^+^ current of RBCs in the mouse retina. We isolated the Ca^2+^-dependent component of the K^+^ current in the mouse rod bipolar cell, and revealed that the IK channel is the origin of the Ca^2+^-dependent K^+^ current. Additionally, we aimed to explore the role of IK in the depolarization and repolarization processes of RBCs and discussed why IK is important for scotopic signal transmission from the RBCs to the AII amacrine cells.

## Results

### 1. Isolation of the Ca^2+^-activated K^+^ currents in the rod bipolar cell

To perform patch-clamp recording from RBCs in retinal vertical slices, we selected the cell bodies of the RBCs at the top of the inner nuclear layer (INL), and their morphological characteristics were visualized by intracellular perfusion of Alexa 488 dye after the whole-cell state (Fig. 1A-B). Rod bipolar cells were stimulated using a step-voltage protocol that stimulates the cells for 2000 ms from a -70 mV holding potential to the target voltage, ranging from -50 to 30 mV at 10 mV intervals. Therefore, the voltage-dependent outward currents were recorded. These outward currents mainly consisted of K^+^ because the Cl^-^-induced outward currents were less than 100 pA at 30 mV, as shown in our previous reports (Paik et al., 2020). In the current traces, we defined a fast component (red arrow), indicating the peak amplitudes between 0 and 200 ms during stimulation, and a sustained component (green arrow), indicating the amplitudes at 25 ms before the end of stimulation (Fig. 1F). Both components of K^+^ currents conspicuously increased from -20 mV, and thus, we hypothesized that they are related to the Ca^2+^ influx from VGCC, reaching a peak around -20 mV in the rod bipolar cell (Protti & Llano, 1998). To confirm the Ca^2+^dependence of the K^+^ current, 5 mM BAPTA was added to the intracellular solution. After a 5-min intracellular approach of BAPTA, the sustained component was significantly reduced to 62% (Fig. 1C-E, n = 5 for control and BAPTA, p<0.05), whereas the fast component was not affected by BAPTA (Fig. 1C-E, n = 5 for control and BAPTA, p>0.05). To evaluate whether the K^+^ current is induced by Ca^2+^ influx through the VGCC, we applied 15 μM mibefradil and 15 μM nifedipine, respectively or concurrently, through the bath application system for 3 min and performed step-voltage stimulation. As shown in Fig. 1F-I, a single application of mibefradil or nifedipine partially inhibited K^+^ currents, whereas the combination of the two inhibitors synergistically inhibited K^+^ currents. In the I-V relationship, the fast components of the K^+^ currents were weakly reduced by a single or concurrent application of mibefradil and nifedipine, while the sustained components were strongly reduced by a single or concurrent application of mibefradil and nifedipine. In particular, mibefradil was found to be more effective than nifedipine was. In details, 35% of the peak amplitudes of the fast components at 30 mV were reduced by mibefradil, nifedipine, and a cocktail of the two inhibitors (n = 7, p>0.05 between the inhibitor groups), while 77% of the sustained components at 30 mV were reduced by mibefradil (with or without nifedipine), and 48% were reduced by nifedipine alone (n = 7, p<0.05, Fig 1J and K). These results suggest that the Ca^2+^-sensitive K^+^ current mainly consists of sustained components of the K^+^ currents and is more strongly related to the mibefradil-sensitive Ca^2+^ channel than the nifedipine-sensitive Ca^2+^ channel is. Based on the effects of the Ca^2+^ channel inhibitors and BAPTA, we determined that the sustained K^+^ current originated from Ca^2+^-activated K^+^ channels.

**Figure 1.**
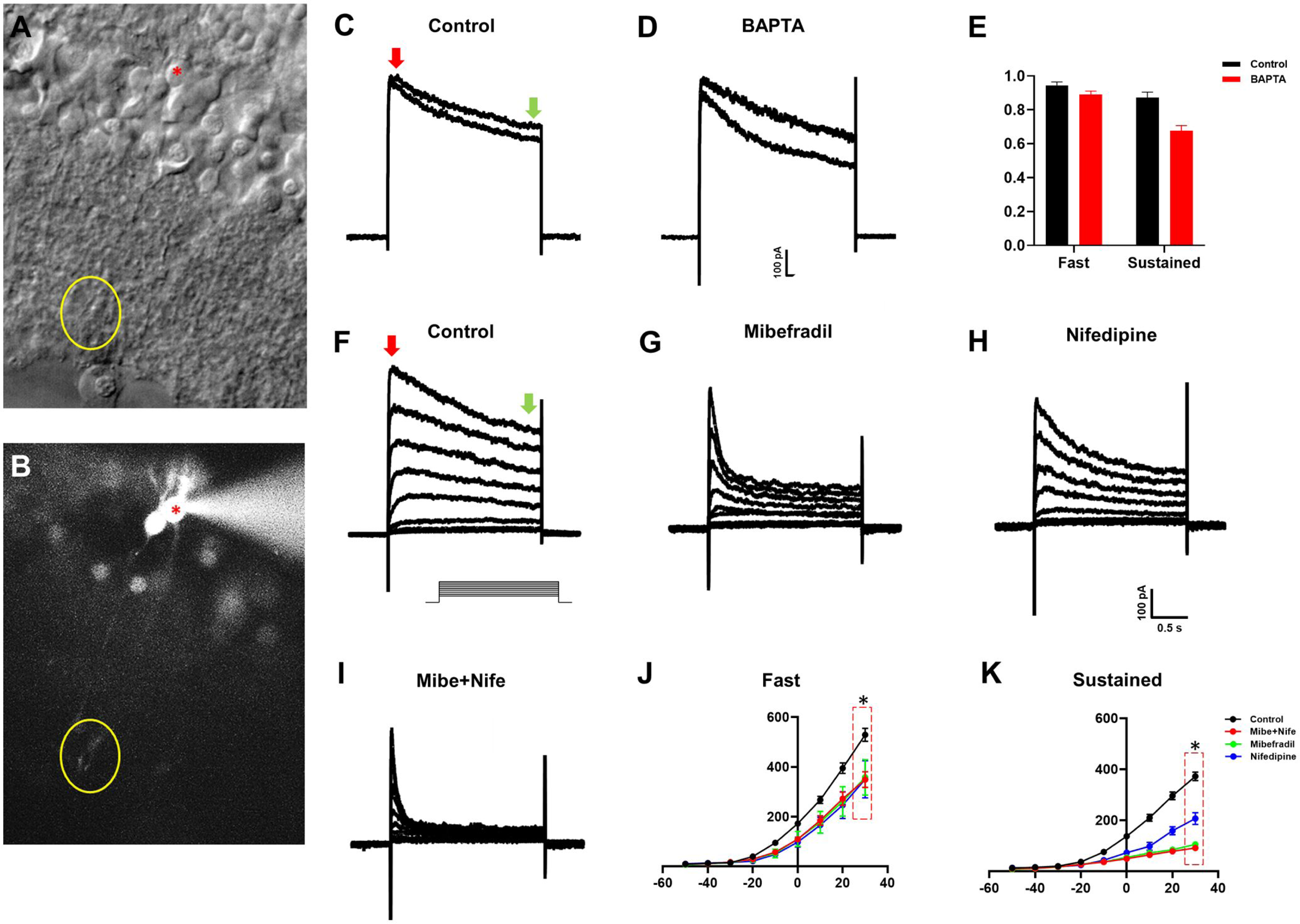
Isolation of the Ca^2+^-activated K^+^ currents in the rod bipolar cells. **A-B.** Selection of the rod bipolar cell in the retinal slice. The cell body of the rod bipolar cell was placed at the top of the inner nuclear layer (asterisk, red), and its axon terminal was detected in the innermost layer of the inner plexiform layer (circle, yellow) through the intracellular diffusion of the Alexa 488 dye. **C-F**. Current traces induced by step-voltage protocol, from -70 mV-holding potential to -50 ∼ 30 mV-target voltage by 10 mV-intervals for 2000 ms, in the control (C), mibefradil (D), nifedipine (E), and mibefradil+nifedipine (F) treated rod bipolar cells. The concentration of mibefradil (a T-type VGCC inhibitor) and nifedipine (a L-type VGCC inhibitor) was 15 μM, and both were pre-applied for 3 min before recording. **G-H.** Current-voltage (I-V) relationship of K^+^ current with the VGCC inhibitors. Fast components were measured at the peak amplitudes in 200 ms from the initiation of the step-voltage (red, 1C), and sustained components were measured at 25 ms before the end of the stimulation (green 1C). In the I-V curve of the fast components (G), outward currents were similarly reduced by VGCC inhibitors (n = 7, p>0.05, ANOVA). In the I-V curve of the sustained components (H), outward currents were strongly reduced by mibefradil or mibefradil+nifedipine than by nifedipine (n = 7, p<0.05 at 30 mV, ANOVA). **I-K**. Effects of the BAPTA on the sustained components. Currents were recorded by step-voltage stimulation from -70 mV to 30 mV. In the control, the currents were minimally changed 5 min after initial stimulation (I), while a definite change of the sustained components was recorded by intracellular perfusion of the BAPTA (J). The differences of the fast components and sustained components were normalized by initial amplitudes and presented by a bar-graph (K). Only sustained components were significantly reduced by BAPTA (n = 5, p<0.05, Student’s t-test).

### 2. Expression of the Ca^2+^-activated K^+^ channels in the rod bipolar cell

To identify the specific types of the Ca^2+^-activated K^+^ channel expressed in the rod bipolar cells, we performed double-labeling immunofluorescence experiments with anti-BK, SK, or IK channel antibodies and anti-PKCα antibody, a specific marker for the rod bipolar cell (Kim et al, 1998; Liets et al, 2006; Negishi et al, 1988). As shown in Fig 2A, BK was mainly detected in the inner segment (IS), OPL, INL, and ganglion cell layer (GCL). SK1 was primarily detected in the proximal portions of the INL and GCL (red in Fig 2B). Additionally, both SK2 (red in Fig. 2C) and SK3 (red in Fig 2D) were detected in the IPL and GCL. However, the immuno-reactivity of anti-BK and SK1-3 antibodies was not co-labeled with the anti-PKCα antibody, which labeled rod bipolar cells (Fig 2A-D). Immunoreactivity of the anti-IK antibody (Alomone) was detected in the IS, OPL, distal and proximal portions of the INL, IPL, and GCL (Fig. 2E). IK immuno-reactivity in the upper portion of the INL was overlapped with PKCα immuno-reactivity, indicating expression of the IK in the cell bodies of the rod bipolar cells (Fig 2E). Taken together, these results suggest that Ca^2+^-activated K^+^ currents in rod bipolar cells, as demonstrated in Fig 1, originate from the IK.

**Figure 2.**
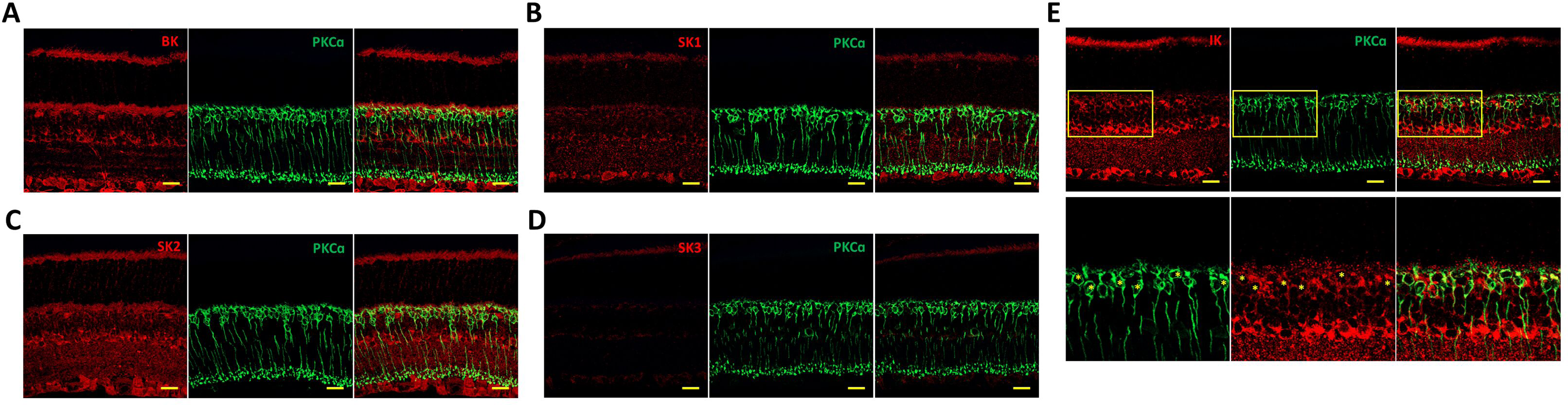
Screening the candidates of the Ca^2+^-activated K^+^ channels by immunohistochemistry. **A.** Co-labeling results of the BK (red) and PKC-α (green) antibodies in the retinal slice. The BK was detected in the inner segment (IS), outer plexiform layer (OPL), upper and lower portions of the inner nuclear layer (INL), and ganglion cells (GC). The BK was not co-localized with PKC-α. **B.** Co-labeling results of the SK1 (red) and PKC-α (green) antibodies in the retinal slice. The SK1 was detected in the lower portion of the INL, ganglion cell (GC), and weakly in the IS. The SK1 was not co-localized with the PKC-α. **C-D.** Co-labeling results of the SK2 (red) or SK3 (red) with PKC-α (green) antibodies in the retinal slice. Both SK2 (C) and SK3 (D) were mainly detected in the GC, and both were not co-localized with PKC-α. **E.** Co-labeling results of the IK (red) and PKC-α (green) antibodies in the retinal slice. The IK (Alomone) was detected in the OPL, upper portion of the INL, IPL, and GC. The IK detected in the upper portion of the INL and innermost layer of the IPL was co-labeled by PKC-α. * Scale bar = 20 μm

### 3. Pharmacological isolation of the IK current in the rod bipolar cell

To confirm whether IK is the origin of Ca^2+^-activated K^+^ currents in rod bipolar cells, we pharmacologically isolated IK currents using the patch-clamp method. We tested the pharmacological effects of iberiotoxin, apamin, and clotrimazole, which are specific BK, SK, and IK channel inhibitors, respectively (Brown et al, 2020; Kshatri et al., 2018). Retinal slices were pre-incubated for 10 min with iberiotoxin (200 nM) or apamin (200 nM), and the step-voltage protocol was applied to the rod bipolar cells to measure changes in K^+^ currents. Compared with the control traces, the K^+^ currents were slightly affected by iberiotoxin and apamin (Fig. 3A-C). In the I-V relationship, the peak amplitudes of the fast and sustained components were minimally affected by iberiotoxin and apamin (n = 6 in each group, Fig. 3D-E). At 30 mV, 6.5% and 8.8% of the fast components were reduced. In addition, 16% and 12.6% of the sustained components were reduced by iberiotoxin and apamin, respectively. We also analyzed the decay time between the 100% and 75% amplitudes to analyze current deactivation. The decay times were not significantly altered by iberiotoxin or apamin (p>0.05, Fig. 3F). These results suggest that functional BK and SK channels are not expressed in rod bipolar cells.

**Figure 3.**
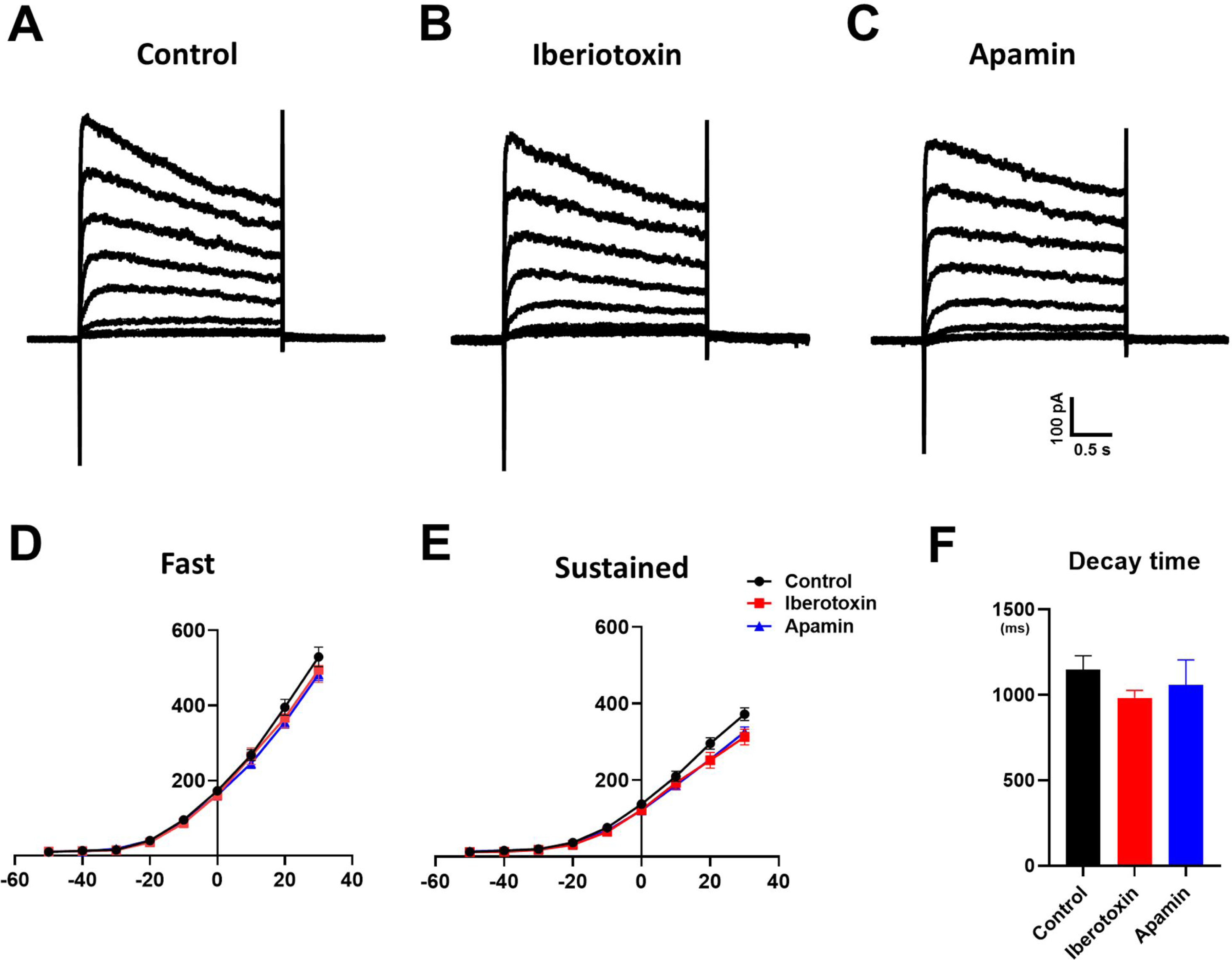
Effects of the BK and SK inhibitors in the rod bipolar cells. **A-C.** The representative traces of the control (A), iberiotoxin (B)-, and apamin (C)-treated groups were presented. Currents were induced by the step-voltage protocol from -70 mV-holding potential to -50 ∼ 30 mV-target voltage. The concentration of the iberiotoxin (BK inhibitor) and apamine (SK inhibitor) was 2 μM, and they were applied for 10 min by bath application system. **D-E.** Current-voltage (I-V) relationship of K^+^ current with the BK and SK inhibitors. Both fast components (D) and sustained components (E) were minimally reduced by iberiotoxin and apamin, and the differences between the two inhibitors were not statistically significant (n = 6, p>0.05 at 30 mV, Student’s test). **F.** Decay time of the current trace at 30 mV. The decay time was measured between 100% (the peak amplitude) and 75% of the amplitudes, and the differences among the control, iberiotoxin, and apamin were not significant (n = 6, p>0.05 at 30 mV, ANOVA).

Next, we used clotrimazole and TRAM34, which are known IK-specific inhibitors. Clotrimazole (15 μM) was pre-treated by bath application for 10 min before achieving whole-cell state, and TRAM34 (1 μM) was applied through the pipette solution for 10 min as described in previous studies (King et al, 2015; Wulff et al, 2001). First, we demonstrated control traces, which were recorded at 0 min after achieving the whole-cell state and 10 min after achieving the whole-cell state (Fig. 4A-B). Current traces changed minimally after 10-minute intervals. Meanwhile, pre-application of clotrimazole effectively inhibited K^+^ currents more than the control current at 0 min did after the whole-cell state (Fig. 4C). The internal application of TRAM34 significantly reduced K^+^ currents at 10 min after the whole-cell state more than the control traces at 10 min after the whole-cell state did (Fig. 4D-E). Almost residual current was inhibited by the addition of 5 μM 4-AP, which is a representative voltage-gated K^+^ channel inhibitor (Fig. 4F). In the I-V relationship, both IK inhibitors successfully inhibited sustained components rather than fast components (Fig. 4G-H). In details, at 30 mV, 22.6% and 10.5% of the fast component were reduced, while 44.2% and 37.5% of the sustained component were reduced by clotrimazole and TRAM34, respectively (n=8, p<0.05, multiple comparison test). The consistent inhibitory effects of both IK inhibitors strongly suggest that the IK channel induces slow and long-lasting outward K^+^ currents in rod bipolar cells. A significant decrease in the decay time between the 100% and 75% amplitudes by IK inhibitors also suggests that the IK channel mediates the sustained component of the K^+^ current (Fig. 4I). Through the inhibitory effect of the IK inhibitors, we clearly demonstrated that IK is a major source of K^+^ current in the rod bipolar cell, specifically inducing sustained components.

**Figure 4.**
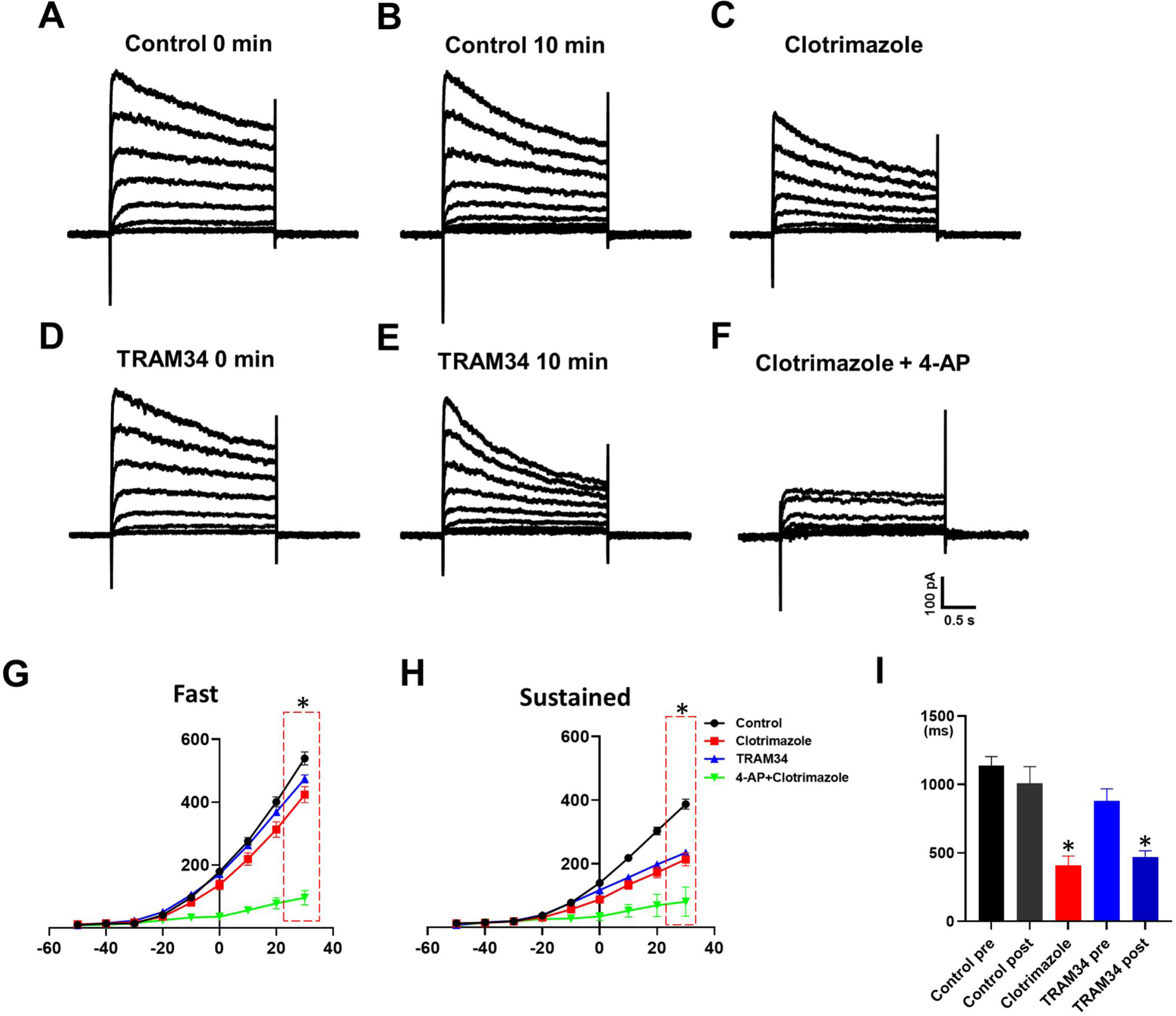
Effect of the IK inhibitors (clotrimazole and TRAM34) in the rod bipolar cells. Clotrimazole (15 μM) was pre-treated by bath application for 10 min, and TRAM34 (1 μM) was applied through the pipette solution for 5 min. Rod bipolar cells were stimulated using a step-voltage protocol from a -70 mV holding potential to a -50–30 mV target voltage. **A-E.** Representative current traces of the control, clotrimazole, and TRAM34-treated rod bipolar cells. Control traces at 0 min (A) and 10 min (B) after whole-cell, clotrimazole-treated traces (C), and TRAM34-treated traces at 0 min (D) and 10 min (E) after whole-cell were presented. Clotrimazole and TRAM34 specifically inhibited the sustained components in the current traces. **F.** Representative current traces for clotrimazole and 4-Aminopyridine (4-AP). The addition of 4-AP, a voltage-gated K^+^ channel inhibitor, reduced the fast components of the current traces. **G-H.** Current-voltage (I-V) relationship of the K^+^ current with IK inhibitors and 4-AP. The fast components were minimally affected by IK inhibitors. However, they were strongly inhibited by the addition of 4-AP (G). The sustained components were effectively inhibited by IK inhibitors, and the addition of 4-AP showed further inhibitory effects (H) (n = 8, P <0.05 at 30 mV, ANOVA).

### 4. IK channel contributes to sustained repolarization after peak membrane potential in the rod bipolar cell

To understand the functional role of the IK channel during rod bipolar cell excitation, we used a current-clamp method and evaluated the effect of an IK inhibitor during depolarization. The basal membrane potential was fixed to -70 mV, and 65 pA was injected for 200 ms to achieve sufficient depolarization of the rod bipolar cell. In the control traces, the injected current depolarized the membrane potential, which showed a rapid rise to a single excitatory peak and sustained subsequent repolarization (Fig. 5A). A rapid single excitatory peak was induced by the VGCCs (Fig. EV1). After the bath application of the 15 μM clotrimazole, rapid depolarization to the single peak from the baseline was not significantly changed compared to control traces, while the repolarization after the single peak was significantly reduced (Fig. 5B). For a detailed analysis, we measured four parameters during the excitation process of the rod bipolar cells, including the peak and plateau membrane potentials from the baseline (-70 mV), after-hyperpolarization, and the recovery time from the end of the current injection to the baseline (Fig. 5A-B). The peak membrane potential and after-hyperpolarization were not statistically different between the control and clotrimazole-treated groups (n = 14, p>0.05, Fig. 5C and E). Whereas the plateau membrane potential was significantly increased by clotrimazole (n = 14, p<0.05, Fig. 5D), and the recovery time from the end of the current injection to baseline was significantly increased by clotrimazole (n = 14, p<0.05, Fig. 5F). The results for the four parameters suggest that IK has little effect on the initial onset of depolarization and mainly controls sustained repolarization, and that the results were temporally matched with the sustained components in the voltage-clamp mode.

**Figure 5.**
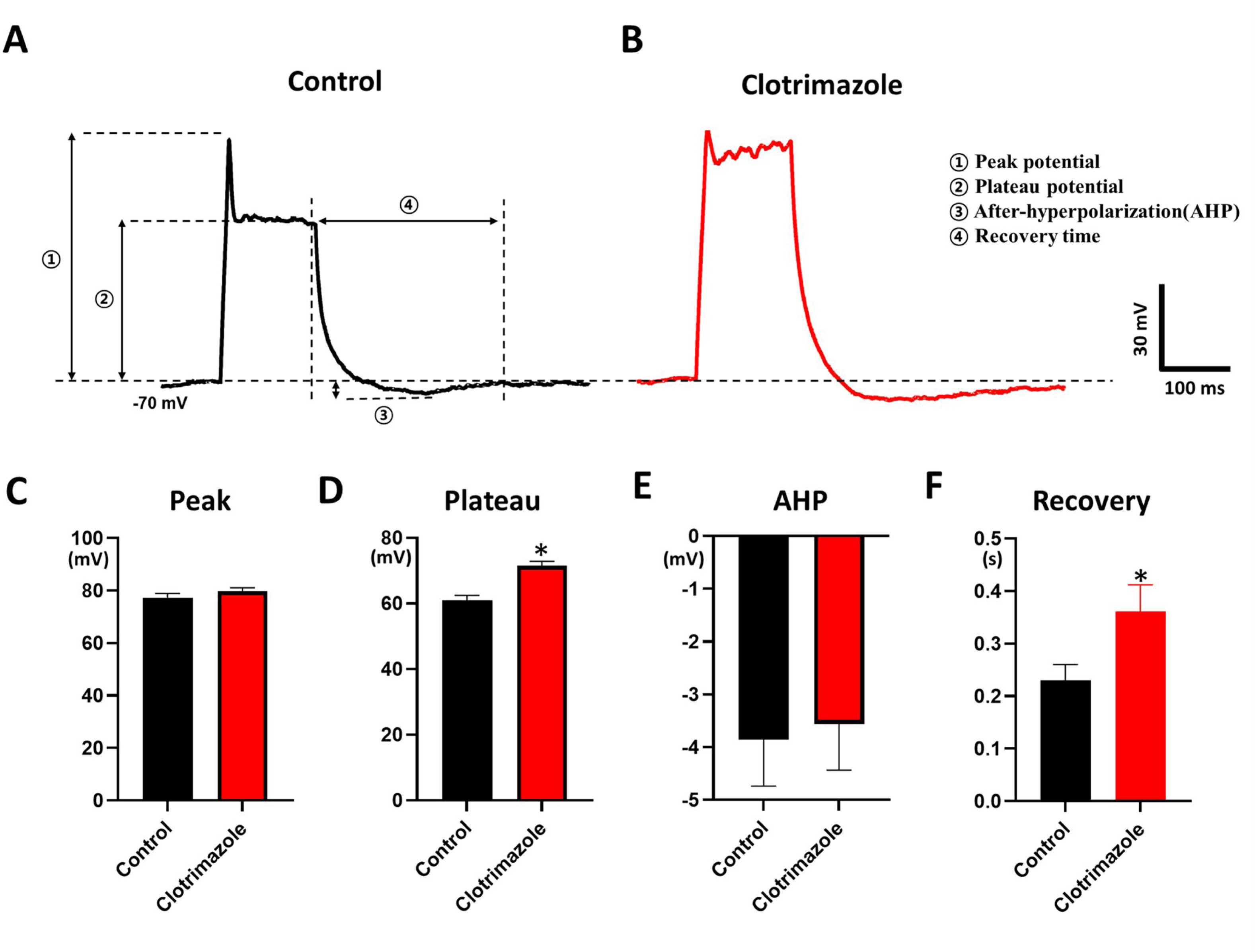
Effect of IK clotrimazole during the rod bipolar cell excitation process in the current-clamp mode. **A-B.** The baseline potential was fixed to -70 mV, and 65 pA was injected for 200 ms. Compared to the control traces (A), repolarization after the early peak was inhibited by the clotrimazole treatment (B). **C-F.** Peak potential (1, C), plateau potential (2, D), after-hyperpolarization (AHP) (3, E), and recovery time from the end of current injection to the baseline (4, F) were analyzed. The plateau potential and recovery time were statistically significantly increased by clotrimazole treatment (n = 14, p<0.05, Student’s t-test), while peak potential and AHP were not changed (n = 14, p<0.05, Student’s t-test).

### 5. IK channel elongates the EPSP duration in synaptic transmission between rod bipolar cell and AII amacrine cell

Inhibition of repolarization by clotrimazole strongly suggests that IK may be involved in synaptic transmission between rod bipolar and AII amacrine cells. To elucidate this, we recorded pairs of rod bipolar cells and AII amacrine cells to record the post-synaptic current from the AII amacrine cells when the rod bipolar cell was stimulated. The cell body of the AII amacrine cells was located at the border of the INL and IPL and had a round- or oval-shaped cell body with an apical single dendrite. After 488 dye perfusion, a narrow field and bistratified morphology throughout the IPL were visualized (Margaret et al, 2019). Rod bipolar cells were selected based on the width of the dendritic field in AII amacrine cells (Fig. EV2). First, we stimulated the rod bipolar cell with a 65 pA current injection to achieve sufficient depolarization and recorded the synaptic current of the AII amacrine cells with or without clotrimazole. If the pair of the rod bipolar cell and AII amacrine cell was selected, fast activation and decay of the synaptic current synchronized with peak excitation of the rod bipolar cells were recorded from the AII amacrine cells (Wan & Heidelberger, 2011). With the application of clotrimazole, repolarization of the rod bipolar cells diminished, and the post-synaptic currents from AII amacrine cells showed elongation of the decay process to the baseline (Fig. 6A, red arrow). To obtain detailed information, we analyzed four parameters of the post-synaptic current: onset time (base to peak), peak amplitudes, decay time (peak to baseline), and area under the curve (AUC; Fig. 6A’). The onset time was significantly reduced, but the difference was only 0.002 s (n = 11, p<0.05; Fig. 6B). The difference in peak amplitudes was not statistically significant (n = 11, p>0.05; Fig. 6C), whereas the decay time and area of the current trace were significantly increased by clotrimazole (n = 11, p>0.05, Fig. 6D and E).

**Figure 6.**
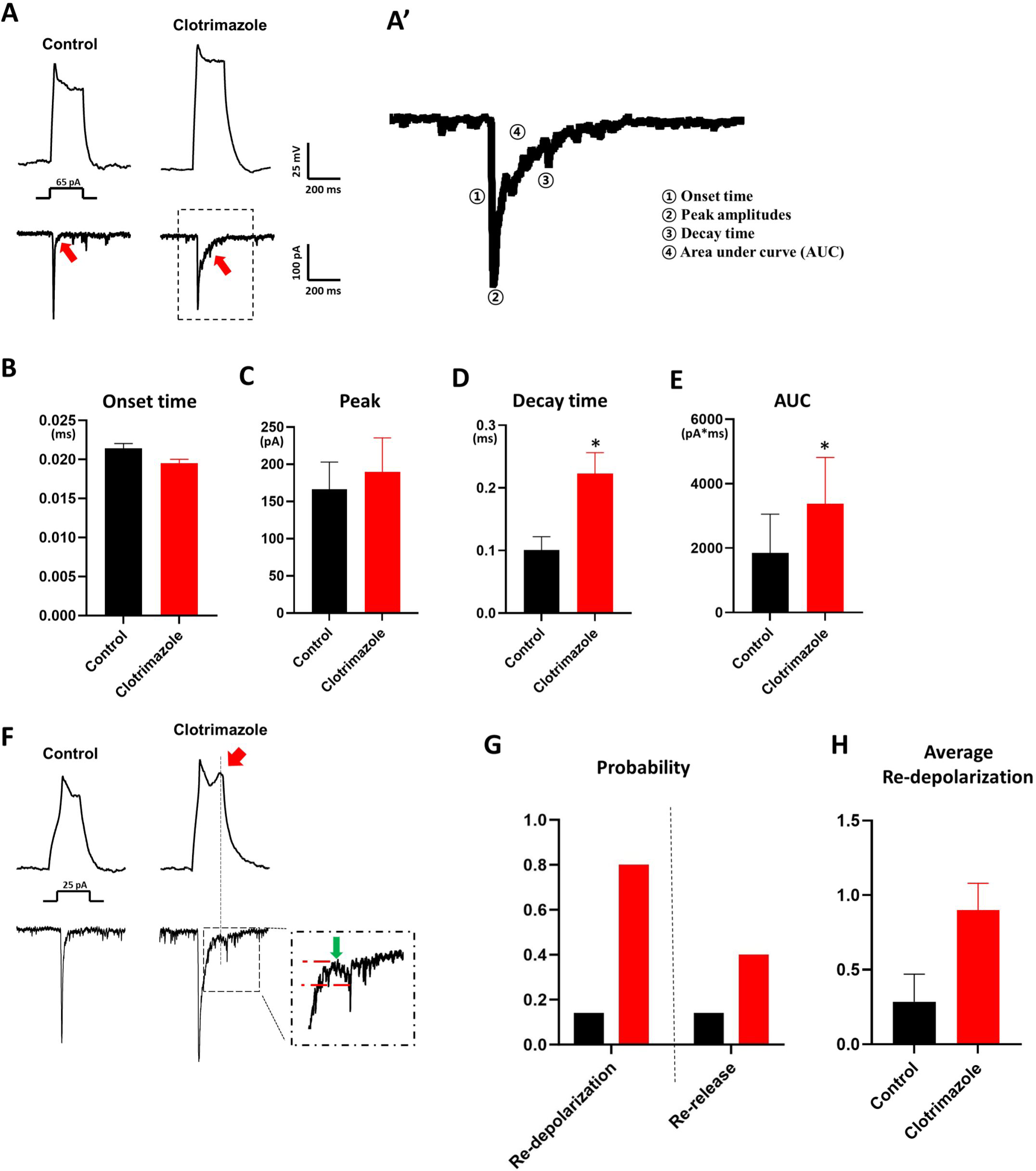
Contribution of the IK in the synaptic transmission between the rod bipolar and AII amacrine cells. **A.** Rod bipolar cells were stimulated by 65 pA injection in the current-clamp mode and synaptic currents were recorded in the AII amacrine cells with -70 mV-holding potential. By the clotrimazole application, repolarization was reduced, and synaptic current was elongated in the AII amacrine cells. **B-E.** Analysis of the synaptic currents in the AII amacrine cells. We measured the onset time (C), peak amplitudes (D), decay time (from peak to baseline, E), and area under curve (AUC, F) of the synaptic current recorded in AII amacrine cells. The rising time, decay time, and AUC were statistically significantly increased by clotrimazole treatment, and the rising time was decreased by clotrimazole (Student’s t-test, n = 10, p<0.05). **F.** Rod bipolar cells were stimulated by 25 pA injection in the current-clamp mode, and synaptic currents were recorded in the AII amacrine cells with a -70 mV holding potential. By the clotrimazole application, the double peak of the membrane potential was recorded in the rod bipolar cells (red arrow) but not in control. The synaptic current of the AII amacrine cells showed weak evocation of the second synaptic current during the decay process of the primary synaptic current (green arrow) by clotrimazole, and the second event was synchronized with the second peak of the rod bipolar cell membrane potential. **G.** The average number of the ‘additional peak’ of the rod bipolar cells (Total additional peak / Total rod bipolar cell number). An additional peak after the initial peak was rare in the control, while an average of one additional peak was detected by clotrimazole-treated rod bipolar cells (n = 8 for control, n = 10 for clotrimazole group). **H**. The probability of the re-opening of the Ca^2+^ channel (double peak) in the rod bipolar cells (double peak rod bipolar cell / total rod bipolar cell, left) and the re-release of the synaptic currents in the AII amacrine cell (a number of second event in AII amacrine cell / double peak rod bipolar cell). Only 20% of the control rod bipolar cells (black) exhibited re-activation, while clotrimazole-treated cells (red) exhibited 80% of the re-activation (left). And, among the pairs of the re-activated rod bipolar cell and AII amacrine cells, 40% of the AII amacrine cells showed re-evoke of the synaptic currents, synchronized with re-activation of the rod bipolar cells (n = 8 for control, n = 10 for clotrimazole group).

Injection of the 65 pA current could induce strong depolarization of the rod bipolar cells, which sufficiently activates the IK channel and induces neurotransmission. Thus, it is beneficial to detect the role of IK channels in the repolarization process. However, a 65 pA current injection could be outside the physiological range compared to the light response of rod bipolar cells (Euler & Masland, 2000; Pang et al, 2004). Therefore, we stimulated rod bipolar cells with a 25 pA current injection to understand the role of the IK current in the relatively weak excitation state. The control traces in Fig. 6F show rapid onset and repolarization in rod bipolar cells and the synaptic current of the AII amacrine cells by 25 pA current injection, and these traces were similar to those of the 65 pA current injection. With the application of clotrimazole, the second peak (red arrow) of the membrane potential occurred in the rod bipolar cells during the repolarization process after the initial peak, indicating re-depolarization. A re-evoked synaptic current was also detected in AII amacrine cells, which was synchronized with the second peak membrane potential of the rod bipolar cells (red dashed line in the inset of Fig. 6F).

For precise quantitative analysis, we manually counted the number of redepolarized membrane potentials in each rod bipolar cell. In the control, 14% (1/7) of rod bipolar cells showed a second peak, whereas 80% (8/10) of the rod bipolar cells showed a second peak after clotrimazole treatment (Fig. 6G, left). And only 14% (1/7) of rod bipolar cells showed re-release of glutamate, inducing a second synaptic current in AII amacrine cells, whereas 40% (4/10) of the rod bipolar cells showed re-release of glutamate (Fig. 6G, right). And “the total number of the second peak / total number of the cell” in the control and clotrimazole-treated group showed an average of 0.1 and 0.9, respectively (Fig. 6H). These results strongly suggest that the IK channel suppresses unintended excitation and synaptic transmission in rod bipolar cells.

### 6. IK channel contributes to faster recovery for consecutive synaptic transmissions

In the above results, we demonstrated that IK channel induces the repolarization of rod bipolar cells, which mediates transient synaptic transmission from rod bipolar cells to AII amacrine cells. These results were obtained after a single stimulation of rod bipolar cells. Thus, we were curious whether the IK channel could affect the consecutive response of rod bipolar cells by repetitive stimulation because cells could prepare for the next depolarization during repolarization. To evaluate this, we designed a two-pulse protocol that stimulates rod bipolar cells twice at 10, 60, 110, 160, 210, 260, and 310 ms intervals and recorded the membrane potential of the rod bipolar cell with the synaptic current of AII amacrine cells. Traces of the two-pulse protocol are presented in Fig. 7A (control) and 7B (clotrimazole). We noticed that the recovery of the peak membrane potential of the rod bipolar cells was slightly altered by clotrimazole. Meanwhile, post-synaptic currents in AII amacrine cells, induced by the second stimulation of rod bipolar cells, were strongly reduced by clotrimazole application (Fig. 7A-B, red arrows). For quantitative analysis, we followed the time change of the peak membrane potential of the rod bipolar cell and the post-synaptic current of the AII amacrine cell in the two-pulse protocol (n = 4). We normalized the second membrane potential and second post-synaptic current using the first membrane potential and first post-synaptic current, as shown in Fig. 7C (membrane potential) and 7D (post-synaptic current). In the control, the second membrane potential was reduced to 86% by a second pulse with a 10-ms interval, and it recovered to more than 97% after a 110-ms interval. Recovery of the membrane potential showed little difference between the control and clotrimazole application groups (p>0.05 at each time point). However, unlike the membrane potential of rod bipolar cells, the post-synaptic currents from AII amacrine cells were significantly altered by clotrimazole. In the control, amplitude of the post-synaptic currents induced by second stimulation at 10 ms was average 11% compared to first stimulation, and it recovered to 27% at 110 ms. With the clotrimazole, amplitude of the post-synaptic currents induced by second stimulation at 10 ms was average 1 % compared to first stimulation, and it recovered to 5 % at 110 ms, which showed statistically lower recovery than the control did (p<0.05, multiple t-test). These results showed that activation of the IK channel had little effect on the peak membrane potential of rod bipolar cells, while it strongly reduced synaptic transmission by repetitive stimulation, which means that IK is important for synaptic function.

**Figure 7.**
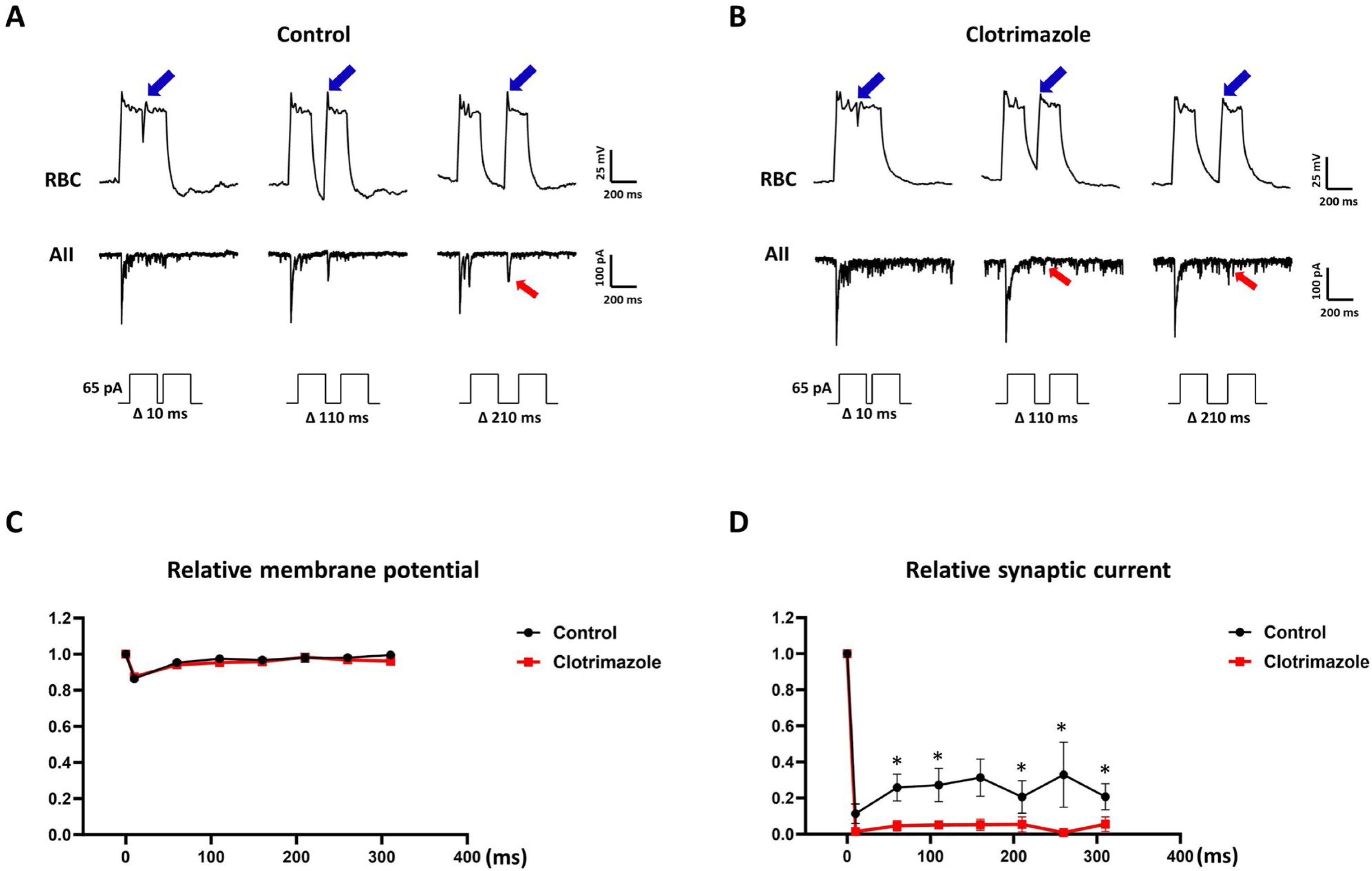
Importance of the IK for consecutive synaptic transmission. **A-B.** Rod bipolar cells were stimulated by 65 pA injection for two times with 10, 110, and 210 ms interval in the current-clamp mode. The synaptic currents were recorded in the AII amacrine cells with a -70 mV holding potential. In the rod bipolar cells, the secondary peak membrane potential (blue arrow) reduced at 10 ms interval, and it recovered similarly with the first stimulation from 110 ms, in both the control (A, upper traces) and clotrimazole application (B, upper traces) traces. Recovery of the secondary post-synaptic current (red arrow) compared to first synaptic current from AII amacrine cells was more reduced in the clotrimazole application group (B, lower traces) than in the control group (A, lower traces). **C.** Time course of the secondary peak membrane potential. We presented the secondary peak membrane by relative values to the first peak membrane potential of the rod bipolar cells. Both the control (black line) and clotrimazole-treated groups showed a decrease at the 10 ms interval pulse and recovered around 95% from the 100 ms interval. **D.** Time course of the secondary synaptic current of the AII amcrine cells. Here, we present the secondary synaptic current relative to the synaptic current. Both the control (black line) and clotrimazole-treated groups showed a decrease at the 10 ms interval pulse. However, after the 60 ms interval, the control group showed more recovery of the secondary synaptic current than the clotrimazole-treated group did. (n = 4, * p <0.05, multiple t-test).

## Discussion

Evaluation of K^+^ channels in neurons is essential for understanding how they control neuronal excitation and synaptic transmission (Goldberg et al, 2005; Trussell & Roberts, 2008). Voltage, Ca^2+^, or other intracellular factors can modulate the gating of various K^+^ channels, which modulate neuronal excitation differently than inhibitory synaptic input like GABA because of different acting times, cellular locations, and gating mechanisms. Thus, various types of K^+^ channels, including voltage-gated K^+^ channel, Ca^2+^-activated K^+^ channel, A-type K^+^ channel, inward rectifying K^+^ channels, have been extensively studied in CNS neurons (Trimmer, 2015). However, although the retina belongs to the CNS, K^+^ channels in retinal neurons are not well understood. Thus, the characterization of K^+^ channels in the main excitatory retinal neurons is needed. A rod bipolar cell is a single-on-type bipolar cell in the scotopic pathway, sharing the same excitatory mechanism as a cone-on-bipolar cell, and it conveys the scotopic signal to the photopic pathway via AII amacrine cells (Marc et al, 2014). Thus, the configuration of the K^+^ channel in the rod bipolar cell could help understand the on-cone bipolar cell excitatory mechanism, which is essential for understanding visual transmission.

In this study, rod bipolar cells showed prominent voltage-dependent outward currents (500–600 pA) in the step-voltage protocol. Additionally, the K^+^ current abruptly increased from -20 mV (Fig. 1C), which is similar to the I-V relationship of VGCCs in rod bipolar cells (Protti & Llano, 1998), which is around -20 mV (Fig. 1C). Thus, we hypothesized that Ca^2+^-activated K^+^ channels are the major components of K^+^ currents. The consistent inhibitory effects of the VGCC inhibitors (mibefradil and nifedipine) and BAPTA on the sustained component of K^+^ currents directly showed Ca^2+^-activated K^+^ currents. The immunohistochemistry results showing IK expression in rod bipolar cells supported our hypothesis (Fig 1-2). We verified the inhibitory effect of IK-specific inhibitors (clotrimazole and TRAM34) on K^+^ currents in rod bipolar cells, whereas BK and SK channel inhibitors were not effective (Fig 3-4). Thus, we could conclude that the IK channel is the source of the Ca^2+^-activated K^+^ currents in rod bipolar cells, based on the negative results of the BK and SK channels of rodent retinal rod bipolar cells in previous studies (Paik et al., 2020; Tanimoto et al., 2012), and the pharmacological definitions indicating that IK is sensitive to IK inhibitors, unlike BK and SK inhibitors (Brown et al., 2020; de-Allie et al, 1996; Ishii et al, 1997).

Previous studies on the IK channel in the CNS mainly focused on non-neuronal cells, including microglia, astrocytes, and endothelial cells (Chen et al, 2015; Hannah et al, 2011; Kshatri et al., 2018; Nguyen et al, 2017; Wulff et al, 2000). Descriptions of IK in CNS neurons have only been reported in a few studies on pyramidal neurons and Purkinje cells. These studies revealed that IK induces after-hyperpolarization (Engbers Jordan et al, 2012; King et al., 2015; Roshchin et al, 2020). However, the role of the IK channels in neural excitation remains unclear. For example, there have been conflicting results regarding after-hyperpolarization in pyramidal neurons (Wang et al, 2016), and the role of IK in synaptic transmission has not been elucidated. This indicates that the detailed role of the IK channel may differ in each type of CNS neuron. In the retina, only Pelucchi et al. revealed that IK is localized in the rod photoreceptor of the salamander retina (Pelucchi et al., 2008). However, this report could not explain the gating mechanism of IK and its functional role because they only isolated the IK current at a membrane potential higher than the physiological range. Therefore, we focused on the gating factor of the IK channel and its contribution to the depolarization process and synaptic transmission in rod bipolar cells. We analyzed four parameters during the depolarization of rod bipolar cells: peak membrane potential, plateau potential, after-hyperpolarization, and decay time in the current-clamp mode. Among them, the plateau potential and decay time were significantly increased by IK inhibitor, indicating that IK is important for repolarization from immediately after peak depolarization to the baseline (Fig. 5). The repolarization effect of IK overlapped with that of ANO1 observed in our previous studies (Jeon et al., 2013; Paik et al., 2020). Although both IK and ANO1 induce Ca^2+^-activated inhibitory signals, IK may have different roles. IK is more sensitive to mibefradil (Fig. 1), whereas ANO1 is similarly affected by mibefradil and nifedipine (Paik et al., 2020). This means that T-type Ca^2+^ channels are very close to IK as a Ca^2+^ source, which is a distinctive feature of the IK channel in rod bipolar cells compared to BK and SK, which form complexes with VGCCs (Berkefeld et al, 2006; Vivas et al, 2017).

Generally, T-type Ca^2+^ channels are low-voltage-activated Ca^2+^ channels that induce low-threshold exocytosis in neurons (He et al, 2018; Weiss & Zamponi, 2013). This has also been reported in retinal rod bipolar cells (Hu et al, 2009; Pan, 2000; Pan et al, 2001). These properties of T-type Ca^2+^ channels may be important for sensing and conveying low-intensity light in the scotopic pathway. In our study, the IK channel was linked to T-type Ca^2+^ channels and induced repolarization. Thus, IK is an essential inhibitory regulator of T-type Ca^2+^ channel-induced low-threshold depolarization in rod bipolar cells, which might result in precise synaptic transmission under dim-light conditions.

To understand the specific role of IK during synaptic transmission, we used a dual patch-clamp method that records pairs of rod bipolar cells and AII amacrine cells. With the 65 pA current injection into rod bipolar cells, we obtained a stable post-synaptic current in the AII amacrine cells. In this protocol, we found that the inhibition of IK in rod bipolar cells elongated the post-synaptic current in AII amacrine cells, while peak amplitudes were not significantly changed (Fig. 6A-E). This indicates that the IK channel plays a role in inhibiting excessive synaptic transmission from rod bipolar cells to AII amacrine cells. Notably, the effect of the IK channel on synaptic transmission between rod bipolar cells and AII amacrine cells differed from that of the BK channel in the photoreceptor terminals in a previous study (Xu & Slaughter, 2005). They reported that a sufficient efflux of K^+^ by BK in the axon terminal of the rod could increase the extracellular K^+^ concentration and promote synaptic release. These conflicting results may be due to the conductance of K^+^ efflux and the expression site of Ca^2+^-activated K^+^ channels. Because IK channels in rod bipolar cells have smaller conductance than BK channels have and are mainly expressed in the cell body, the IK channel cannot sufficiently increase the extracellular K^+^ concentration compared to the BK channel in the photoreceptor axon terminal.

To evaluate the function of IK in a more physiological range, we reduced the amplitude of the current injection (25 pA) to rod bipolar cells to analyze the role of IK in rod bipolar cell excitability. Low amplitudes of current injection reflect low-threshold VGCC activation, which might be helpful for mimicking dim light stimulation in the rod pathway. With the 25 pA current injection with IK inhibition, rod bipolar cells showed a double-peak pattern during depolarization, which was not detected in the control state (Fig. 6). Because activation of the VGCC is a unique source of the fast and transient peak of the membrane potential in the rod bipolar cell (Fig. EV1), the double peak suggests the re-activation of VGCCs. The re-activation of the VGCCs could be interpreted in two ways: the re-opening of the VGCCs after deactivation and the activation of the remaining VGCC, which was not recruited for the first peak. Although we could not know the exact reason for the double-peak, it is certain that the second depolarization occurred by insufficient repolarization due to IK inhibition. After fast depolarization and repolarization, the membrane voltage decreased to below the threshold of the VGCCs; thus, the VGCCs could remain in a resting state. However, due to IK channel inhibition, insufficient repolarization maintains the membrane voltage of the rod bipolar cells over the threshold of the VGCC; thus, VGCCs could be in a re-activation state. This indicates that the IK channel inhibits the over-excitation of rod bipolar cells for precise synaptic transmission.

Repolarization is a critical step in neural excitation because the gating of the K^+^ channel sufficiently reduces the membrane potential to close the voltage-gated ion channels; thus, excitatory voltage-gated cation channels can prepare for the next excitation (Cooper, 2012; Grider et al, 2023). In this study, we revealed that the IK channel plays a role in the repolarization of rod bipolar cells, suggesting that the IK channel plays a role in the consecutive excitation of rod bipolar cells and synaptic transmission. To evaluate this, we performed a two-pulse protocol and analyzed the secondary membrane potential of rod bipolar cells and the synaptic currents of AII amacrine cells. Interestingly, the IK channel had little effect on the recovery of the peak membrane potential of the second stimulation, whereas inhibition of the IK channel reduced the secondary synaptic current (Fig. 6). These results indicate that the IK channel suppresses excessive synaptic transmission in a single stimulation, which is important for preparing the next excitation of the rod bipolar cell and the AII amacrine cell pathway.

In summary, we demonstrated the distinct characteristics of Ca^2+^-activated K^+^ channels in rod bipolar cells. The IK channel is activated by a low-threshold Ca^2+^ channel, which induces a sustained component of K^+^ currents and affects the repolarization process. Due to sustained K^+^ currents, the IK channel specifically modulates the repolarization process rather than the peak membrane potential of rod bipolar cells. This may be due to the distinctive role of the IK channel compared to the voltage-gated K^+^ channels in rod bipolar cells. In synapses, IK plays a complex role. First, it suppresses excessive prolongation of the synaptic current. Second, IK suppress the unintended re-activation of Ca^2+^ channels in rod bipolar cells during the repolarization state. Third, IK is essential for consecutive activation of rod bipolar cells via the AII amacrine cell pathway.

We isolated a unique type of K^+^ channel, IK, from rod bipolar cells. There is little known about the role of the IK channel in the CNS, including retinal neurons, and the inhibitory mechanism of rod bipolar cells is not well known. Therefore, we expect that our results will expand our understanding of the role of IK in the neural system as well as visual transmission. However, we still need further evaluation of IK channels in the retina because our immunoreactivity showed IK expression in amacrine cells, and IK expression in the other bipolar cell subtypes is unknown. Moreover, understanding the role of IK in macroscopic retinal function is required to completely understand the contribution of IK to vision.

## Method

### 1. Ethical standards

All animal experiments followed the regulations of the Catholic Ethics Committee of the Catholic University of Korea, based on the guidelines of the National Institute of Health (NIH) for the Care and Use of Laboratory Animals (NIH Publications No. 80-23). The animal experimental protocol was approved by the Institutional Animal Care and Use Committee and Department of Laboratory Animals of the College of Medicine, The Catholic University of Korea (Approval Number: CUMC 2020-0245-06).

### 2. Retinal slice preparation

C57BL/6 mice (6-7 weeks old) were used for patch-clamp experiments. The mice were kept under a 12-hour light/dark cycle in a climate-controlled laboratory. As described in our previous study (Paik et al., 2020), acute retinal slices were prepared to perform patch-clamp method. For slice preparation, we used oxygenated, with 95% O_2_–5% CO_2_ gas, iced extracellular solution (NaCl 126, KCl 2.5, CaCl_2_ 2.4, MgCl_2_ 1.2, NaH_2_PO_4_ 1.2, NaHCO_3_ 18, and glucose 11, in mM). After anesthetization by intraperitoneal injection of zolazepam (20 mg/kg) and xylazine (7.5 mg/kg), the eyes were enucleated, and the mice were sacrificed by an overdose of zolazepam. The anterior compartments of the eye, including the cornea and lens, were immediately removed from the iced extracellular solution, and the retina was gently dissected from the sclera. Retina was embedded in 1.5% agar and sectioned into 200 μm thickness using a vibratome (Campden instrument, Loughborough, England). The prepared retinal sections were incubated in the oxygenated extracellular solution at room temperature for 1 h (24–27℃).

### 3. Visualization of the rod bipolar cell and AII amacrine cell

The retinal slices were observed using a BX51 microscope (Olympus, Tokyo, Japan). The cell bodies of rod bipolar cells are located at the top of the INL, adjacent to the OPL. After achieving the whole-cell state, rod bipolar cells were confirmed by their morphological characteristics through the intracellular diffusion of the Alexa 488 dye, which showed a single projecting axon to the innermost layer of the IPL, with two or three axon terminals. The cell bodies of the AII amacrine cells were located at the border between the INL and IPL with a thick apical dendrite projecting into the IPL. AII amacrine cells were confirmed by their morphological characteristics after perfusion with Alexa 488 dye, whose processes were bistratified in the IPL by lobular appendages in sublamina *a* and stratifying dendrites in stratum five (Veruki & Hartveit, 2002; Zandt et al, 2017).

### 4. Electrophysiology

A recording solution containing NaCl, KCl, MgCl2, CaCl2, HEPES, and Glucose (in mM) was prepared, and its pH was adjusted to 7.4 with NaOH. A pipette solution containing KCl 30, K-gluconate 100, MgCl_2_ 1, EGTA 0.5, ATP-Na, ATP-Mg, and GTP-Na (in mM) was prepared, and its pH was adjusted to 7.4 with KOH. A MultiClamp 700 B amplifier and Axon Digidata 1440A (Molecular Devices, San Jose, CA, USA) were used for voltage- and current-clamp recording. Pipette resistance was 10 to 12 MΩ in the extracellular solution when it was filled with pipette solutions. The recording pipettes were tightly attached to the cell membrane to achieve a gigaohm seal. After the giga-ohm seal state, the whole-cell state was achieved using a brief application of negative pressure. The pipette resistance and whole-cell capacitance were compensated by the MultiClamp 700 B panel. The stimulation protocols in the voltage-clamp and current-clamp methods were designed using pClamp 10 software (Molecular Devices).

### 5. Immunohistochemistry

For the immunohistochemistry, eyecups were fixed with 4% PFA for 1 h, and they were kept in 30% sucrose in PB 0.1M overnight to prepare frozen blocks using the OCT compound. The frozen tissues were sectioned by 8 μ m thickness using the LEICA cryostat (LEICA, Wetzlar, Germany). Retinal sections were washed with 0.01 M PBS and boiled with 10 mM citrate buffer for antigen retrieval. After then, sections were blocked by a cocktail of 10% normal donkey serum, 2% bovine serum albumin, and 0.2% Triton X-100 for 1 h, and those were incubated with primary antibodies for 18 h at 4°C. The sections were washed in PBS and incubated with a secondary antibody for 2 h at room temperature. Anti-KCa1.1 (1:100; Alomone, Jerusalem, Israel), Anti-KCa2.1 (1:100; Alomone), Anti-KCa2.2 (1:100; Alomone), Anti-KCa2.3 (1:100; Alomone), Anti-KCa3.1 (1:100; Alomone), Anti-PKCα (1:1000; Santa Cruz, Dallas, Texas, U.S.A.), Cy3 (1:1500; Jackson Immunoresearch, West Grove, PA, USA), and Alexa 488-conjugated antibodies (1:500; Molecular Probes, Eugene, OR, USA) were used for primary and secondary antibodies. All images were captured using a Zeiss LSM 900 confocal microscope (Carl Zeiss Co. Ltd., Oberkochen, Germany).

### 6. Data analysis

The amplitude and decay time of the patch-clamp results were measured using the Clampfit software (Molecular Devices). Statistical analyses, including the Student’s t-test and one-way ANOVA, were performed using the Prism 8.0 software (GraphPad, San Diego, CA, USA) to determine statistical significance, with a p-value of < 0.05.

## Acknowledgement

This research was supported by the Basic Science Research Program through the National Research Foundation (NRF) of Korea, funded by the Ministry of Education (grant number 2021R1I1A1A01049783 [YSP]) and by the Basic Science Research Program of the National Research Foundation (NRF) of Korea, funded by the Ministry of Education, Science, and Technology (NRF-2022R1A2C2006178 [I-BK]).

## Author contribution

Yong Soo Park

Funding acquisition, electrophysiology and immunohistochemistry, and original draft writing

Ki-Wug Sung

Supervision, data analysis, review, and editing

In-Beom Kim

Conceptualization, funding acquisition, supervision, data analysis, review, and editing

## Supplementary

**Figure EV1.**
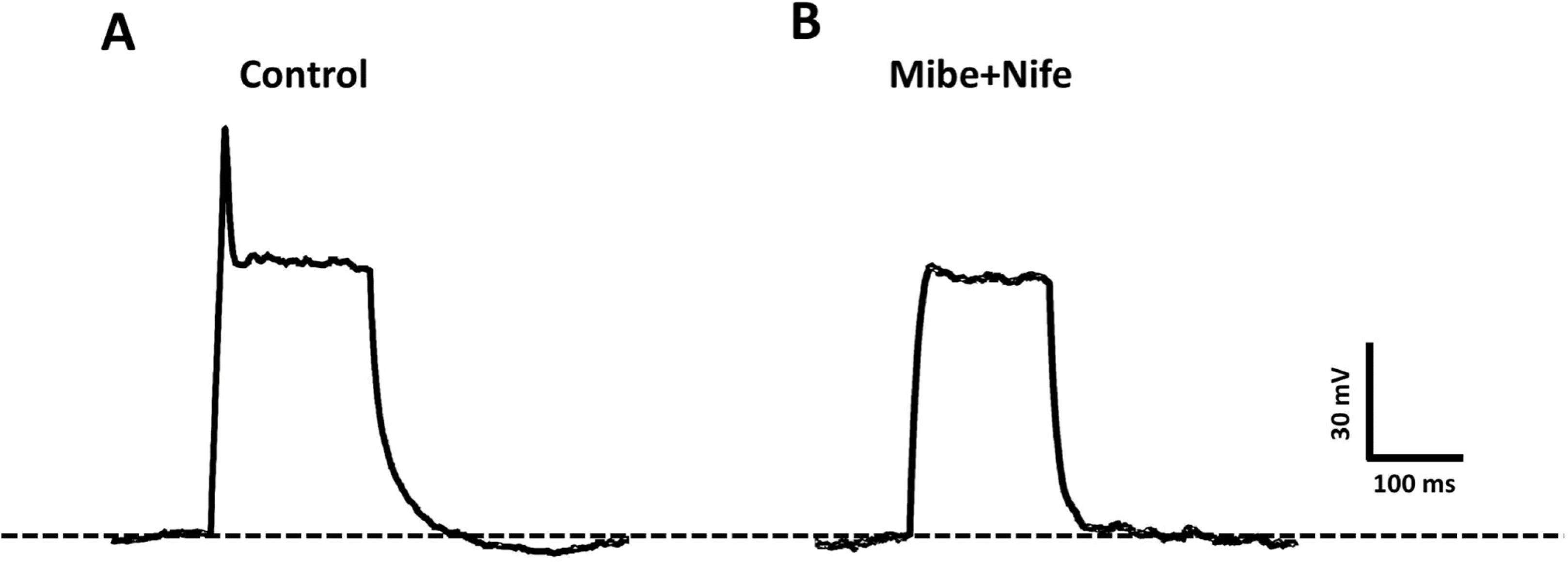
Calcium-dependent excitation of the rod bipolar cells. **A.** Excitation traces of the rod bipolar cells. Rod bipolar cells were stimulated by a 65 pA current injection. **B.** Effect of the Ca^2+^ channel inhibitors on rod bipolar cell excitation. The combination of 15 μM mibefradil and 15 μM nifedipine completely abolished fast excitatory peak membrane potential.

**Figure EV2.**
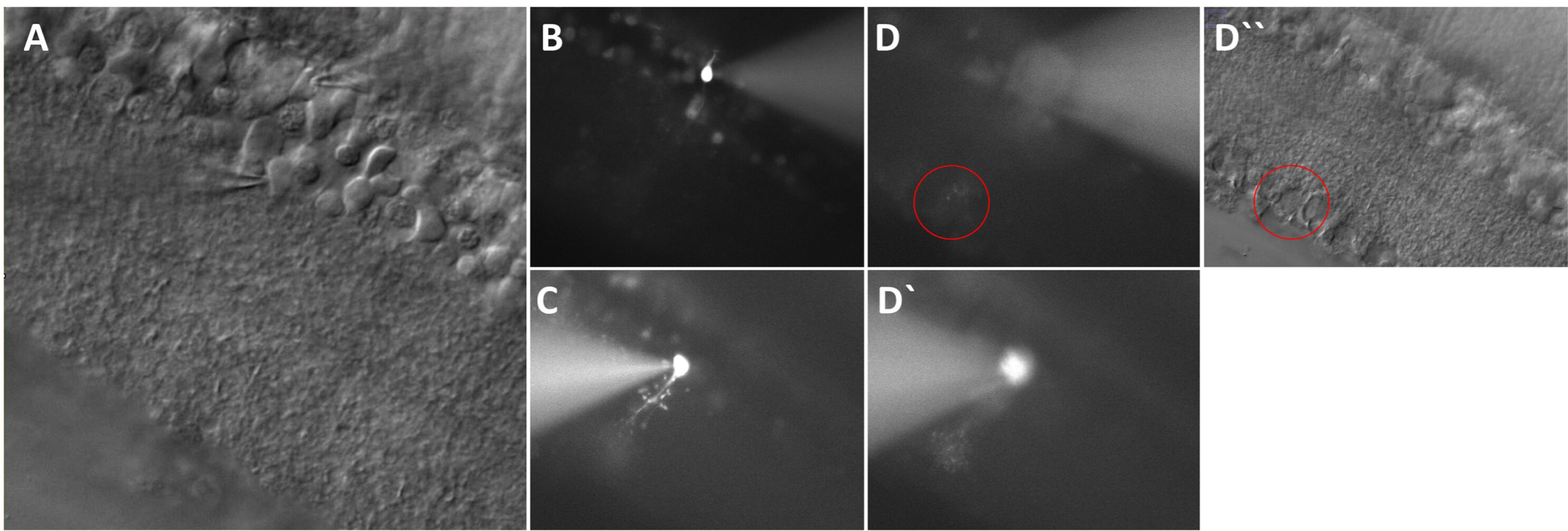
Selecting the pair of the rod bipolar cell and AII amacrine cell. **A.** Selection of the pair of the rod bipolar cell and AII amacrine cell in the DIC image of the retinal slice. The cell body of the rod bipolar cell was selected in the top of the INL, and the cell body of the AII amacrine cell was selected by two characteristics. One is inverted cone-shaped with apical dendrite, and the other is half of the cell body buried in the IPL. **B.** After the perfusion of the Alexa 488 dye, rod bipolar cell showed 2∼3 axon terminals located in the innermost layer of the IPL. **C.** After the perfusion of the Alexa 488 dye, AII amacrine cells showed bistratified processes in the On/Off layer of the IPL with appendage in the proximal to cell body. **D**. Contact of the axon terminal of the rod bipolar cell and AII amacrine cell dendrite in the IPL.

